# Functional and metagenomic level diversities of human gut symbiont-derived glycolipids

**DOI:** 10.1101/2023.05.23.541633

**Authors:** Ji-Sun Yoo, Byoungsook Goh, Kyoo Heo, Da-Jung Jung, Wen Zheng, ChangWon C. Lee, Naama Geva-Zatorsky, Meng Wu, Seung Bum Park, Dennis L. Kasper, Sungwhan F. Oh

## Abstract

Bioactive metabolites produced by symbiotic microbiota causally impact host health and disease, nonetheless, incomplete functional annotation of genes as well as complexities and dynamic nature of microbiota make understanding species-level contribution in production and actions difficult. Alpha-galactosylceramides produced by *Bacteroides fragilis* (BfaGC) are one of the first modulators of colonic immune development, but biosynthetic pathways and the significance of the single species in the symbiont community still remained elusive.

To address these questions at the microbiota level, we have investigated the lipidomic profiles of prominent gut symbionts and the metagenome-level landscape of responsible gene signatures in the human gut. We first elucidated the chemical diversity of sphingolipid biosynthesis pathways of major bacterial species. In addition to commonly shared ceramide backbone synthases showing two distinct intermediates, alpha-galactosyltransferase (agcT), the necessary and sufficient component for BfaGC production and host colonic type I natural killer T (NKT) cell regulation by *B. fragilis,* was characterized by forward-genetics based targeted metabolomic screenings. Phylogenetic analysis of agcT in human gut symbionts revealed that only a few ceramide producers have agcT and hence can produce aGCs, on the other hand, structurally conserved homologues of agcT are widely distributed among species lacking ceramides. Among them, alpha-glucosyl-diacylglycerol(aGlcDAG)-producing glycosyltransferases with conserved GT4-GT1 domains are one of the most prominent homologs in gut microbiota, represented by *Enterococcus bgsB*. Of interest, aGlcDAGs produced by bgsB can antagonize BfaGC-mediated activation of NKT cells, showing the opposite, lipid structure-specific actions to regulate host immune responses. Further metagenomic analysis of multiple human cohorts uncovered that the *agcT* gene signature is almost exclusively contributed by *B. fragilis*, regardless of age, geographical and health status, where the *bgsB* signature is contributed by >100 species, of which abundance of individual microbes is highly variable. Our results collectively showcase the diversities of gut microbiota producing biologically relevant metabolites in multiple layers-biosynthetic pathways, host immunomodulatory functions and microbiome-level landscapes in the host.

## Introduction

Gut microbiota is a unique ecosystem with unmatched genetic, molecular, and species-level diversities. The metagenome of GI symbionts outnumbers the host genome by hundreds to thousands of folds, capable of producing small molecules (metabolites) with enormous structural variety(Skelly et al., 2019). Since a significant portion of metagenome remained unannotated or uncharacterized, identification of novel molecular structures does not always come with a clear explanation of biosynthetic mechanisms. Redundancy at gene and pathway levels among many bacterial species makes this question more complex to solve at the microbiota level. Furthermore, gut microbiota undergoes continuous and dynamic changes over time; such variability makes efforts to determine the biological relevance in the host-microbiota context difficult.

Alpha-galactosylceramides (aGCs) from gut microbes are one of the first symbiont-derived metabolites with characteristic host immune activities(An et al., 2014). Originally found in the marine sponge and opportunistic pathogens as immunostimulatory agonists of NKT cells, ubiquitous human gut symbiont *B. fragilis* produces structurally distinct aGCs (BfaGCs) with immunomodulatory functions. BfaGCs regulate the early proliferation of NKT cells in the colon, as well as induce immunomodulatory effector function of NKT cells, of which structure-activity relationship was shown among their natural structural variants(Oh et al., 2021). Nonetheless, essential genes for BfaGC (and for the bacterial sphingolipids in general) biosynthesis are only partially characterized in gut microbiota. Furthermore, it remained unclear whether B. fragilis is the only BfaGC producer in gut microbiome, or whether other species with conserved biochemical machinery can also produce aGCs or structurally related and bioactive glycolipid species.

To systematically address these questions, we have developed a multi-pronged approach. Firstly, sphingolipid biosynthesis profiles of the major gut symbionts were acquired and responsible genes including unique alpha-galactosylceramides (*agcT*) were characterized. Aa expanded search with alpha-glycosyltransferase domain identified an alpha-glucosyldiacylglycerol synthase, which are widely distributed among human infant gut symbionts. Immunological assessment of bacterial lipids was carried out to confirm host actions, followed by gene signature analysis of human microbiota cohorts of different age, ethnicity and disease groups.

## Results

### Biochemical diversities of core gut symbionts in sphingolipid production

Accumulating reports from multiple research groups show several classes of sphingolipids of gut bacterial origin, which can contribute to host physiology in several aspects (An et al., 2014; Brown et al., 2019; Johnson et al., 2020; Le et al., 2022; Oh et al., 2021). Nonetheless, the sphingolipid profile and biosynthetic capability of prominent gut symbionts are only partially elucidated (An et al., 2011). We investigated >1200 gut bacterial species (obtained from multiple human gut metagenomic datasets (Pasolli et al., 2017), whose reference genome and protein were all available at NCBI database. More than 600 species had Spt homologs (> 30% sequence identity and < 10-50 e-value) in their individual genomes. Among these species, we selected 12 major species in the human gut and profiled sphingolipids of individual bacterial cultures (Figure 1A). The presence and absence of possible intermediates were unambiguously assigned by MS/MS fingerprints (Figure S1).

**Figure 1.**
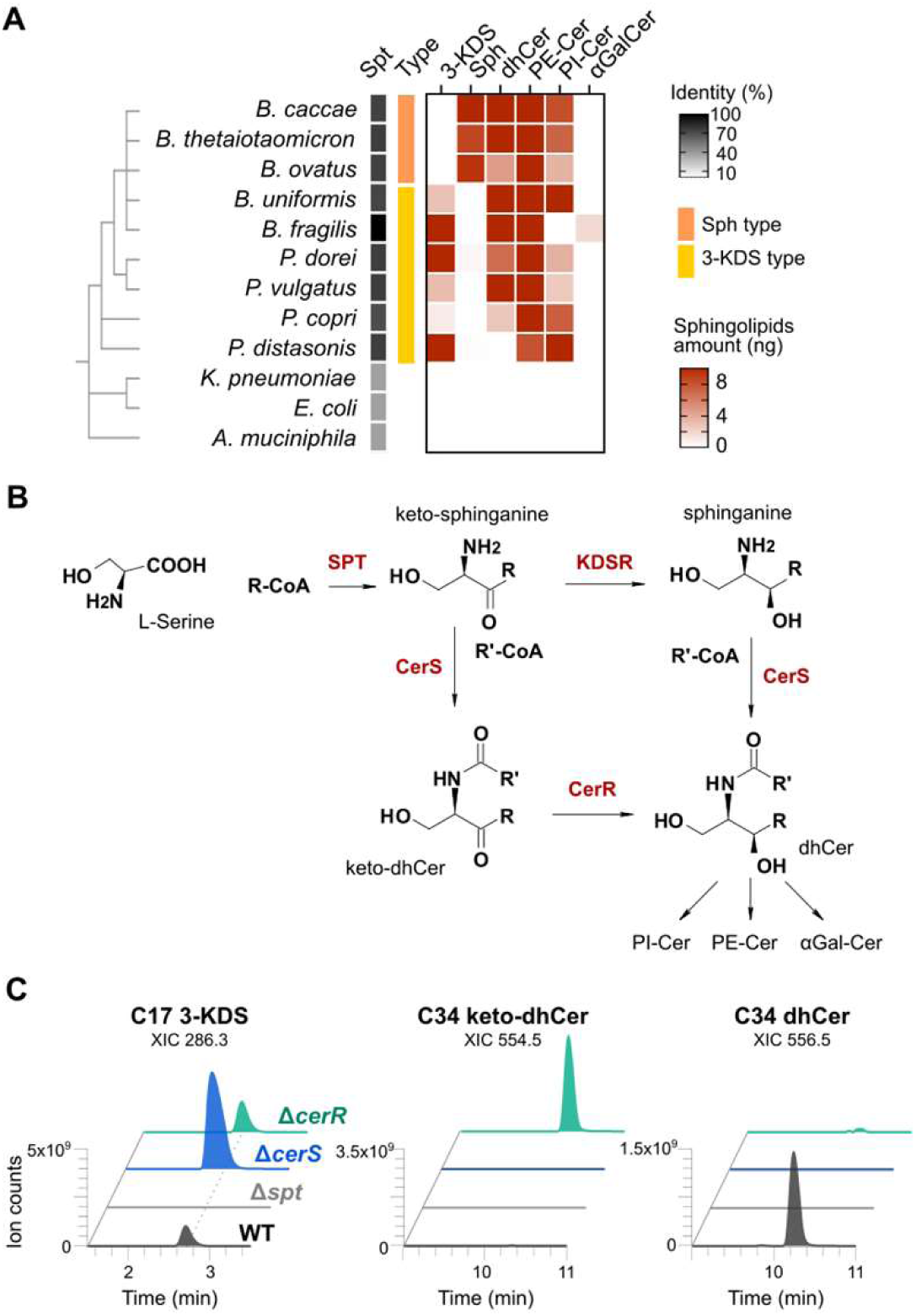
Biochemical diversity of sphingolipid-producing gut symbiont. A) Lipidomic profiles of sphingolipid-producing gut symbiotic Bacteroidales show clear separation in two groups, producing 3-KDS or sphinganine as a major intermediate. B) Two potential pathways of gut bacterial sphingolipid synthesis based on the lipid intermediate profiles. C) Sphingolipid profiles of individual synthase knockout strains of *B. fragilis* confirm 3-keto-dhCer as an intermediate, kinetically controlled by *B. fragilis* CerR.

Of note, lipidomic profiling revealed two distinct pathways of bacterial sphingolipid biosynthesis, confirmed by both sphingolipid intermediates. A group of bacterial species (Figure 1A, color-coded in yellow) including *B. fragilis* has significant levels of 3-ketosphinganine (3-keto-dihydrosphingosine or 3-KDS) as a major intermediate. To further delineate species-specific biochemistry (figure 1b), we investigated responsible genes in *B. fragilis* sphingolipid biosynthesis, from homology search of *Caulobacter crescentus* CCNA_01212 and CCNA_01222[(Stankeviciute et al., 2021). Two *B. fragilis* genes, BF9343_4218 and BF9343_4240 were identified as putative ceramide synthase (CerS) and ceramide reductase (CerR), respectively. We established isogenic knockout mutants of two genes (Figure S3A) and obtained sphingolipid profiles of each mutant. As previously shown, Δ*spt* cannot synthesize any of sphingolipids(An et al., 2014), in comparison, Δ*cerS* results in the accumulation of 3-KDS (Fig S3A-C) and the loss of *cerR* caused the accumulation of 3-keto-4,5-dihydro-ceramide (keto-dhCer) (Fig S3D-G). These results collectively suggest that 3-KDS is the substrate of CerS, producing keto-dhCer as a transient intermediate, which is further reduced by CerR (Figure 1C).

In parallel, other ceramide producers including *B. thetaiotaomicron* and *B. ovatus* (Fig 1A, color-coded in orange), 3-KDS is nearly undetectable, instead its putative reduction product spinganine is identified as a major intermediate. In these species, 3-ketodihydrosphingosine reductase (KDSR) dominantly reduces 3-KDS to sphinganine, which is further conjugated to ceramides by CerS homologues, which resembles known the biosynthetic route with their eukaryotic counterparts (Lee et al., 2022).

### Targeted metabolomic screening identified agcT as the necessary and sufficient gene for BfaGC biosynthesis (Fig 2)

**Figure 2.**
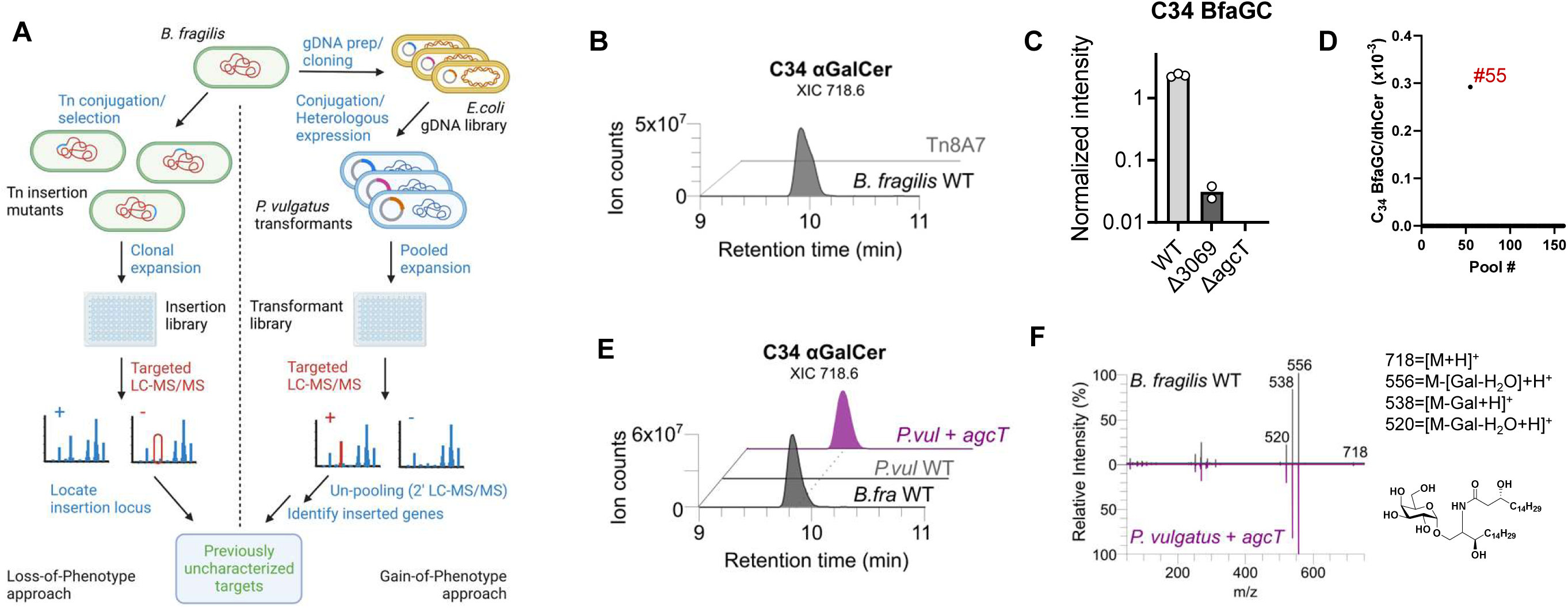
Identification and functional characterization of B. fragilis alpha-galactosyltransferase (agcT) by forward genetic screening approaches. A) Graphical scheme of loss-of-function and gain-of-function library generation and targeted lipidomic analysis. B) Identification of single transposon insertional mutant Tn8A7 showing selective loss of BfaGCs. C) Downstream synthase agcT KO strain shows a total loss of BfaGCs. D) LC-MS-based screening of the B. fragilis genome transformant library uncovered a single pool of transformants, producing BfaGCs. E-F) agcT-transformed *P. vultagus* produces identical molecule to BfaGC shown in XIC (E) and MS/MS spectra mirror plot (F).

In contrast to ceramide synthesis, alpha-hexosyl linkage between sugar and ceramide backbone is unique to bacterial lipids. <u>T</u>o identify the putative alpha-galactosyltransferase in *B. fragilis*, we generated a genome-wide, clonally isolated transposon mutant library (Goodman et al., 2009)and subsequently performed targeted lipidomic screening (Figure 1A). From more than 5000 mutants analyzed, one transposon mutant (Tn8A7) exhibited a deficiency in BfaGC production, while ceramide biosynthesis remained unaffected (Figure 1B and S1A). The insertion site of the Tn8A7 genome was identified as BF9343_3069 locus. An isogenic knockout mutant (ΔBF9343_3069) were generated and assessed ceramide production of the mutant. ΔBF9343_3069 produced comparable amounts of dihydroceramide (dhCer) and dihydroceramide phosphoethanolamine (PECer), but the synthesis of BfaGCs is diminished by ∼98% (Figure S1B). Based on protein motifs and predicted functions (putative two-component sensor kinase response regulator), BF9343_3069 is hypothesized to function as an upstream regulator of BfaGC synthesis rather than an authentic alpha-galactosyltransferase. To further identify the downstream effector enzyme involved in the synthesis of BfaGCs, we conducted transcriptomic analysis and found 400 differentially expressed genes in ΔBF9343_3069 compared to the wild type. Among BF9343_3069 regulon (Figure S2A), 20 putative glycosyltransferases were identified to be significantly downregulated. Only one of the candidate glycosyltransferases, BF9343_3149, does not belong to capsular polysaccharide operon and has the glycosyltransferase GT4 domain, which is involved in the transfer of glycosyl group with retention of the anomeric carbon stereochemistry thus can produce alpha-hexosyl linkage. (Comstock et al., 1999; Coyne et al., 2008; Moremen & Haltiwanger, 2019). We generated an isogenic mutant of BF9343_3149 (Δ*agcT*) (Figure S1C) and confirmed the complete loss of BfaGC production (Figure 1D).

In parallel to loss-of-function screening, we conducted a gain-of-function screening by generating a transformant library of *B. fragilis* genome to confirm the gene function unambiguously (Figure 2A). We introduced this *B. fragilis* genomic DNA library into ceramide-producing *P. vulgatus* that do not produce aGCs. From ∼17500 colonies combined to 160 pools (∼110 clones per pool) we identified a positive pool out of 160, which exhibited the production of aGCs (Fig 2). We isolated single colonies from the positive pool, 2 clones were identified to produce aGCs. The transformed elements were individually sequenced and confirmed as *agcT* gene. We cloned full-length *B. fragili*s agcT and heterologously expressed it in *P. vulgatus* and *B. thetaiotaomicron* (Fig S). The production of aGCs was observed in both species, providing further evidence that AgcT is a sufficient enzyme for the production aGCs from ceramides, consistent with a recent biochemical study of the gene(Okino et al., 2020)

### BfaGC is a necessary and sufficient factor for NKT cell regulation in vitro and in vivo. (Fig 3)

**Figure 3.**
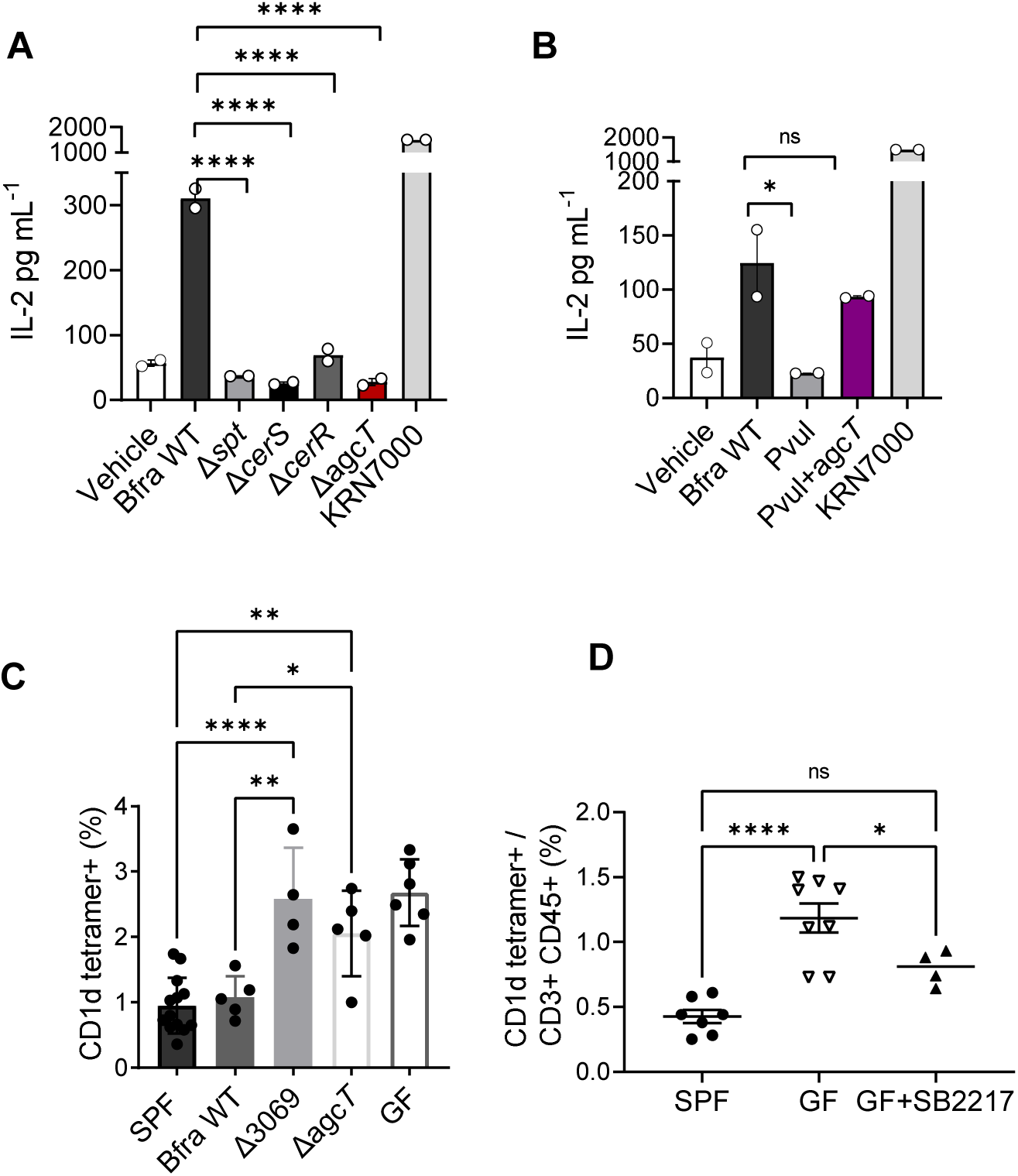
BfaGC is a necessary and sufficient factor to modulate NKT cells. A) Deletion of individual gene in BfaGC biosynthesis loss CD1d-mediated IL-2 inducing activity in NKT cell elicited by B. fragilis total lipids. B) Introduction of agcT to ceramide producer species enables to induce IL-2. C) Selective loss of BfaGC by deletion of agcT or upstream regulator BF9343_3069 abrogate modulatory function of colonic NKT cell development by B. fragilis. D) Oral introduction of synthetic BfaGC (SB2217) to germ-free mice in early life can normalize colonic NKT cells.

Next, we assessed in vitro and in vivo NKT cell modulation of B. fragilis sphingolipids. An APC-free NKT cell assay system was employed (Brennan et al., 2011; Cheng et al., 2006; Naidenko et al., 1999; Zeissig et al., 2013) by pre-incubating murine CD1d with lipid extracts from individual strains. As expected, none of lipid extracts isolated from *B. fragilis* deletion mutants of aGC biosynthesis pathway genes (*Δspt, ΔcerS, ΔcerR, ΔagcT)* cannot elicit IL-2 responses from NKT cells (Figure 3A). On the other hand, lipid extract from *P. vulgatus* which heterologously expresses *agcT* can induce IL-2, showing necessity and sufficiency of the *agcT* (Figure 3B).

We have previously shown that the loss of entire sphingolipid species from *B. fragilis* blunts *in vivo* regulatory impact to colonic NKT cells, which can be rescued by oral supplementation of purified BfaGCs [An 2014]. Monocolonization of BfaGC-deficient mutants (ΔBF9343_3069 or Δ*agcT*) resembles colonic NKT cell dysregulation shown by total sphingolipid KO, unambiguously confirming that BfaGC is the necessary factor. In contrast, germ-free mice given synthetic BfaGCs at neonatal stage show normalized colonic NKT cells, implying that NKT cell regulatory impact by symbiont-derived ligands can happen even in absence of gut microbiota and microbiota-derived immunological niche(Olszak et al., 2012).

### Metagenome-wide sequence similarity search identified BgsB as a prominent structural homolog of B. fragilis AgcT in gut microbiota, producing alpha-glucosyl-diacylglycerols (aGlcDAGs)

With *B. fragilis agcT* characterized its function to synthesize BfaGC in vitro and in vivo, we expanded our investigation other ceramide-producing gut symbionts with capability to synthesize aGCs. BLAST search for *agcT* homologs revealed that among *spt*-positive Bacteroidales, very few (a total of 16 species) possess homologs of *agcT*. Out of these 16 species, 12 species were found to harbor both *cerS* and *cerR* genes as well. We performed targeted sphingolipid analysis on *B. fragilis* strains, which is the most abundant species among *agcT*-positive species in human gut, and *B. salyersiae*, the second most abundant species. Targeted LC-MS/MS analysis confirmed that all *B. fragilis* strains tested, and one of the two *B. salyersiae* strains, exhibited the presence of the aGCs.

As described, AgcT possesses a characteristic protein domain, cd03817 GT4_UGDG-like glucosyltransferase [NCBI Conserved Domain Database, Figure S4]. We hypothesized that gut bacterial species containing cd03817 domain protein in their genome are putative producers of modulators for NKT cells. Based on this domain, we expanded homology search and found that cd03817 family proteins are widely distributed in the human metagenome. Of interest, prevalent species with cd03817 family proteins largely belong to order Lactobacillales (Figure 4A-B) in infant gut, where the symbionts critically contribute to colonic immune development(An et al., 2014; Olszak et al., 2012). Several representative species (*S. mitis, L. rhamnosus* and *E. faecalis*) with cd03817 family proteins were chosen and the production of alpha-glucosyldiacylglycerols (aGlcDAGs) was confirmed (Figure 4C-D and S5A-B).

**Figure 4.**
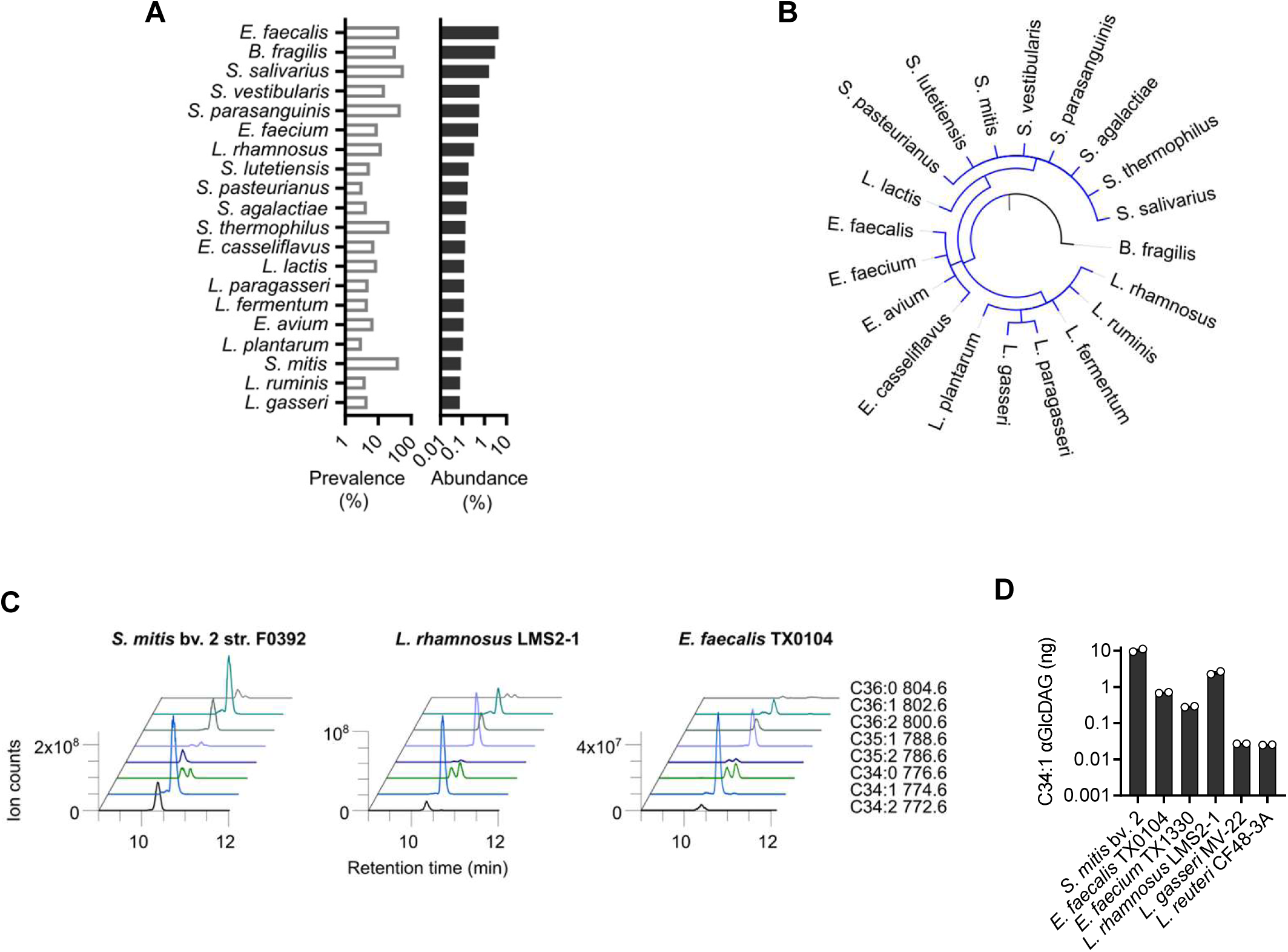
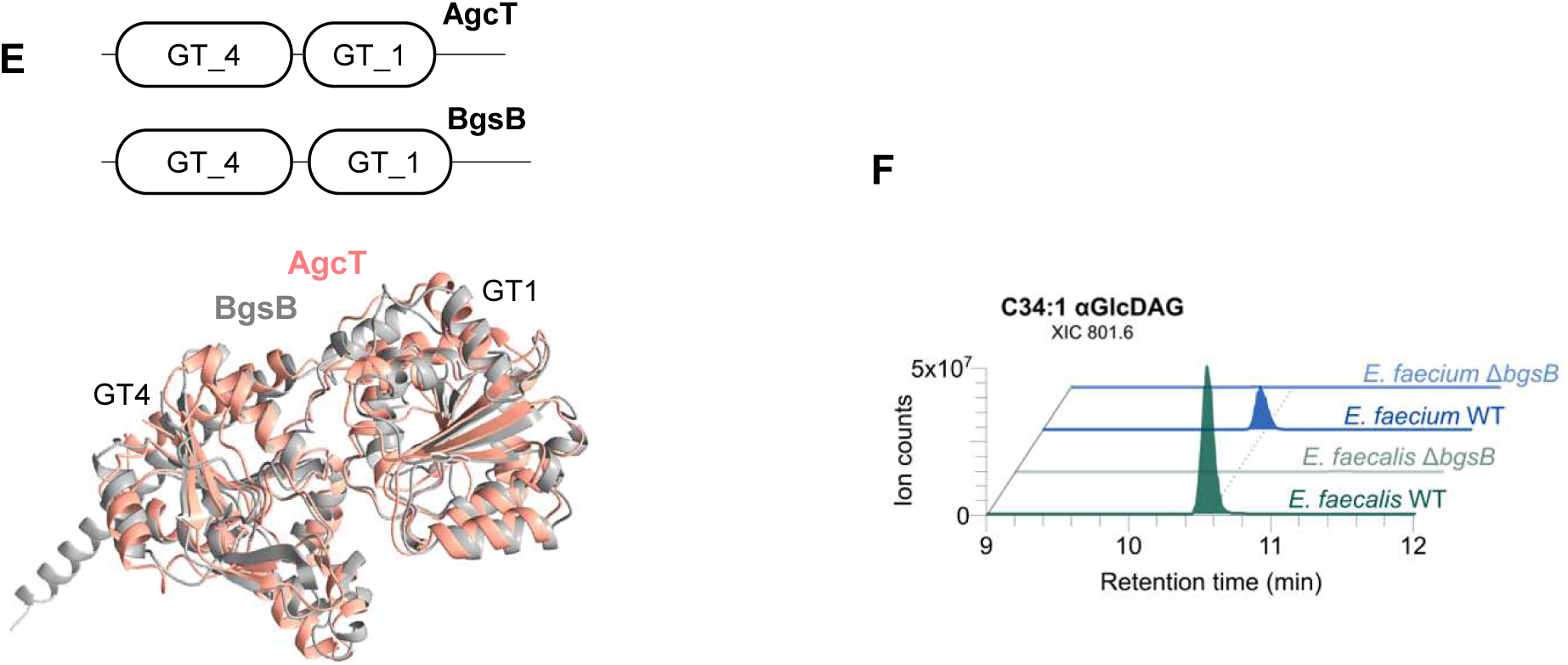
Metagenome-wide ortholog analysis uncovers that B. fragilis agcT is structurally related to bgsB, a widely shared in gut microbiota producing alpha-glucosyldiacylglycerol (aGlcDAG). A) Twenty most prevalent and abundant species in the infant metagenome that harbor the GT4_UGDG-like domain protein. B) 19 out of 20 top species belong to order Lactobacillales except B. fragilis. C) aGlcDAGs profiles of cd-3817 GT4+ species, showing chain-length variation. D) Quantitation of C34:1 aGlcDAG level in cd-3817 GT4+ species. E) Pfam motifs and superimposition of *in silico* predicted 3D structures show structural similarities of AgcT and BgsB, both bearing glycosyltransferase Family 4 (GT4) and glycosyl transferases group 1 (GT1) domains. F) bgsB is a responsible gene for aGlcDAG production in Enterococcus. XIC of C34:1 aGlcDAGs of *E. faecalis* and *E. faecium* WT and bgsB KO mutants.

For further structural and genetic studies, we chose *E. faecalis*, the most abundant cd03817 species in infant metagenome as the target species. *E. faecalis* BgsB is annotated as a glucosyltransferase to produce aGlcDAGs (Theilacker et al., 2009, 2011). Although the sequence identity of two proteins is only 27%, the *in silico* predicted AlphaFold structures of *B. fragilis* AgcT and *E. faecalis* BgsB exhibit a significant conservation (Figure 4E). Various strains of *Enterococcus* produce aGlcDAGs (S5C) and isogenic KO mutants in *E. faecalis* and *E. faecium bgsB* confirmed that the gene is necessary for the aGlcDAG biosynthesis (Fig 4F and S5D).

### aGlcDAGs antagonize BfaGC actions on NKT cells. (Fig 5)

Since some alpha-hexosyl-diacylglycerols from pathogenic bacteria (such as aGal-DAGs (BbGL-2) of *Borrelia burgdoferei* were reported to be NKT agonists (Kinjo et al., 2006), we assessed whether aGlcDAGs produced by *bgsB* homologs can also function as CD1d ligands and NKT cell agonists. To our surprise, aGlcDAGs producing-symbiont lipids did not elicit NKT cell activation (Fig 5A), however antagonize BfaGC-mediated NKT cell activation in dose-dependent manner (Fig 5B). The bacterial lipids isolated from *bsgB* knockout strains can activate nor inhibit NKT cells (Fig 5), confirming their antagonistic activity to NKT cell is aGlcDAG-specific. In parallel, unlike *B. fragilis*, monoassociation of *E. faecalis* at birth did not modulate colonic NKT cell proliferation (Fig 5D).

**Figure 5.**
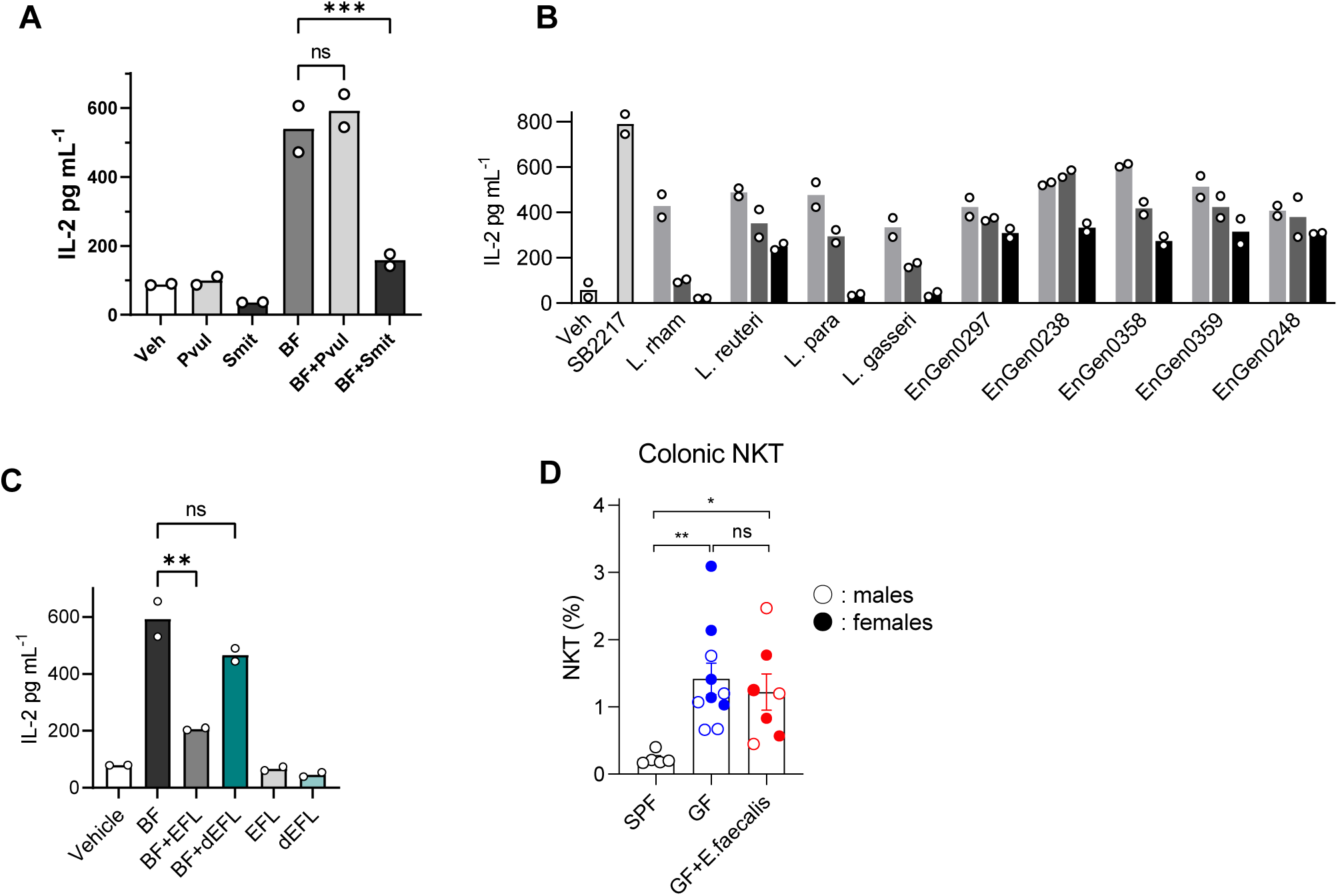
aGlcDAGs antagonize BfaGC actions on NKT cells. A) IL-2 secretion of 24.7 NKT cells incubated with CD1d pre-loaded bacterial lipids from *B. fragilis* with or without lipids from *P. vulgatus* and *S. mitis* in a 1:1 ratio. B) Bacterial lipids from various cd03817+ species exhibit dose-dependent antagonism to SB2217. C) IL-2 secretion of 24.7 NKT cells incubated with CD1d pre-loaded bacterial lipids from *B. fragilis* with or without EFL and ΔbgsB mutant in a 1:4 ratio. D) Monocolonization of E. faecalis cannot normalize colonic NKT cell development.

### agcT and BsgB signatures show distinct patterns over multiple human metagenome cohorts (Fig 6)

Finally, to profile the landscape of these two structurally related but functionally distinct bacterial gene signatures in human gut microbiota, we have investigated abundance of *agcT* and *bsgB* homolog signatures from multiple human metagenome cohorts covering diverse ethnicity, ages and disease status(Pasolli et al., 2017). Most significantly, in longitudinal data of early human microbiota (up to 4 years of age), two homolog groups show distinct abundance patterns. The absolute majority (>95%) of agcT signature is originated from *B. fragilis*, in contrast, contributing species for *bgsB* signature is much more diverse and dynamically changes over the early development (Figure 6). Observed microbiota landscape also holds in subgroups, divided by countries or modes of delivery: *bgsB* signature varies, where *agcT* is almost monopolized by *B. fragilis* (Figure S6B).

**Figure 6.**
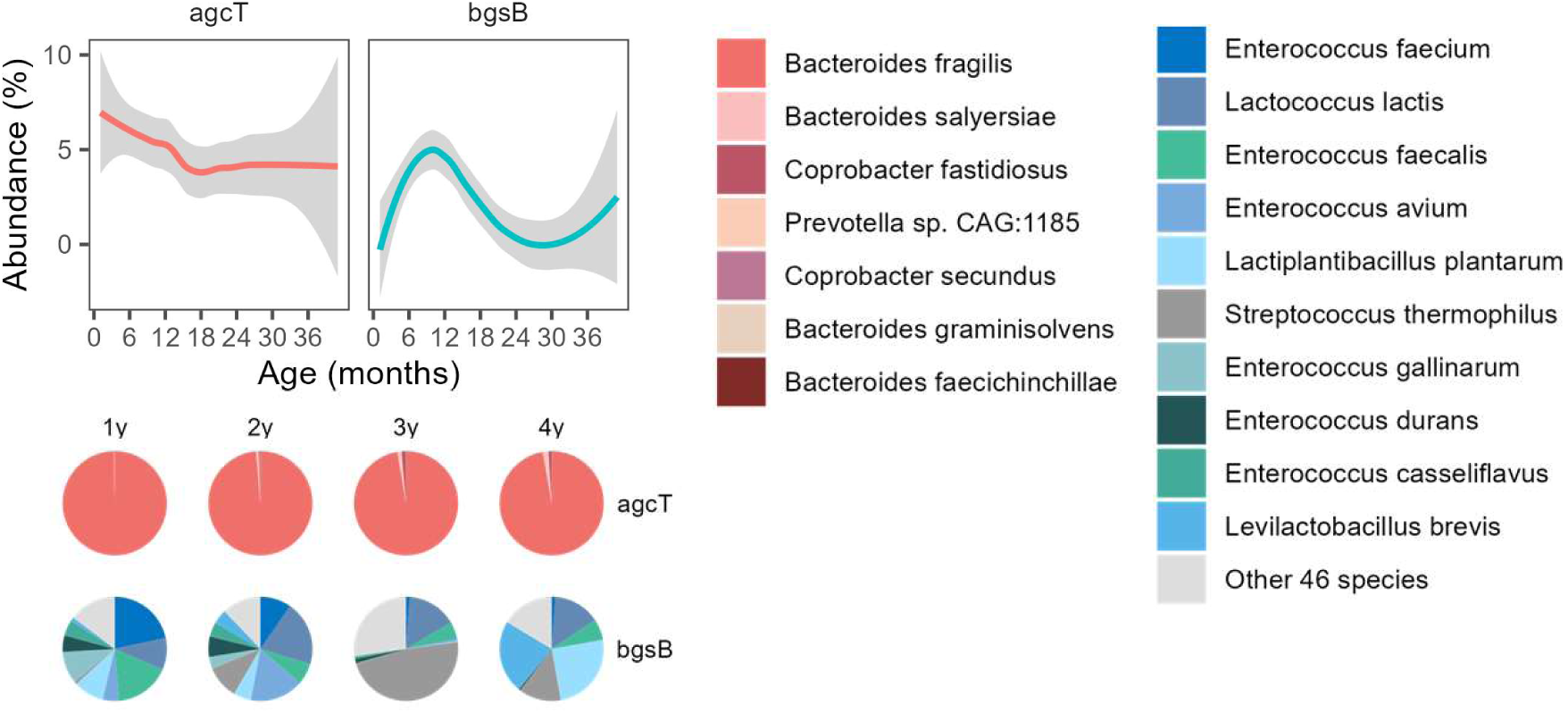
B. fragilis dominated agcT+ abundance, whereas various species constitute bgsB+ signature in infant gut microbiota.

## Discussion

Having co-evolved over millions of years, multicellular hosts have developed sophisticated mechanisms to precisely distinguish innocuous or harmful microbes, based on the recognition of specific structural components. A subtle structural difference can determine the responses from full-blown immune stimulation or complete tolerance. Gut symbiont-derived sphingolipids are recognized as prominent lipid species in the GI lumen, nonetheless, structural diversity originated by different bacterial species and their impact to the host has not been well understood. Our current results show metabolomic diversity of gut symbionts even in the same order, whose major intermediates (3-KDS and sphinganine) may affect host physiology in structure-specific manner (Liu et al., 2022; Spears et al., 2022).

In addition to the potential activities as intermediates, bacterial ceramides are conjugated to structurally and functionally diverse sphingolipids. Non-peptide small molecules of microbial origins are traditionally regarded as primary stimuli of the innate immune system; Pattern recognition receptors (PRR) cover a vastly broad range of molecules with distinct chemical groups, from oligonucleotides to heavily acylated saccharolipids. Recent studies on nonclassical presentation molecules such as MHC class I-like proteins have broadened the recognition of nonpeptide ligands beyond PRRs (Mayassi et al., 2021; Oh et al., 2022). These presentation molecules, such as CD1d or MR1, restrict oligomorphic T cell receptors of unconventional T cells (NKT cells and mucosa-associated invariant T (MAIT) cells, respectively). Considering unconventional T cells are a prominent component of host immunity where constant interaction with residential microbes takes place (such as mucosal layers and skin), microbial contribution to the unconventional T cell function is crucial.

In this juncture, regulation of colonic NKT cells by gut microbiota and identification of aGCs from *B. fragilis* have served as one of the prototypic cases in that symbiont-derived metabolites can exert immunomodulatory functions. BfaGCs are structurally and functionally distinct from aGCs of non-symbiotic (environmental or opportunistic pathogenic) origins and BfaGCs can regulate NKT cell proliferation in early development, as well as the effector function at the adult stage. Nonetheless, even *B. fragilis* is regarded as ubiquitous in human microbiota and highly represented in early human gut(Stewart et al., 2018), it was still unclear the contribution of *B. fragilis* as the single component of the gut microbial community. Here we demonstrate metabolomic and metagenomic profiles of sphingolipid biosynthesis from gut symbionts, not only that *B. fragilis* is the most prominent producer of symbiont-derived aGCs but also that *B. fragilis* is by far the most dominant component among aGC-producing bacteria in human gut microbiota.

The mammalian gastrointestinal tract is home for diverse bacterial species, numbering in the hundreds or even thousands. Many of these species are phylogenetically related and share a large number of homologous genes. However, it is important to note that while these homologous genes may share similarities based on the protein classifications, these similarities do not always translate into the metabolite production and/or recognition. Both AgcT and BgsB are members of the glycosyltransferase family 4 proteins and are classified under the same class (EC number 2.4.1.337) in the enzyme database. As shown, they also share a conserved retaining glycosyltransferase domain and are assigned in the same KEGG orthology (K19002). However, our comprehensive analysis has revealed that despite these similarities, the metagenomic and functional characteristics were distinct. The metagenomic analaysis indicates significant differences in the distribution and prevalence of *agcT* and *bgsB* among the microbial communities. Functionally, agcT is involved in the synthesis of aGCs and mediates NKT cell activation while bgsB is involved in the biosynthesis of aGlcDAGs and can antagonize NKT cell activation by aGCs. These results highlight the diverse and specific functions of glycosyltransferase of glycolipids within microbiota.

We demonstrated that homologous genes in microbiota may exhibit divergent metabolite production profiles and immunological properties. Therefore, it is crucial to consider that while protein clustering based on domain structures provides a useful framework for understanding the function of microbiota, it is essential to delve deeper into the specific metabolic capabilities and functional characteristics of each species to gain a comprehensive understanding of their roles within the complex ecosystem of the mammalian gut. The comparison between *agcT* and *bgsB* signatures in multiple human cohorts further confirmed the diversity in host-microbiota context. Theoretically, two extremes in metabolite formation in the multi-species community can be considered: 1) a single species is entirely responsible for a specific metabolite or 2) all and every species is equally capable of. Our metagenomic analysis implies that the agcT is likely to be close to the former scenario, where the bgsB is the later case. Considering the structural similarity of these two classes, the observation also implies that sequence-and/or structure-based classification of microbiota metagenome requires cross-validation at the molecular product level for proper functional prediction. Along with that our previous observation that BfaGC abundance is linearly correlated with the in human microbiota associated mice supplemented with B. fragilis, the result collectively suggests that B. fragilis is the dominant contributor of BfaGC in the gut. Further metatranscriptome and/or (meta-)metabolome level investigation may elucidate expression and/or specific activity of individual gene products.

### Limitation of Study

Although *B. fragilis* is a prominent member of early gut microbiota, whether human colonic NKT cell development is directly contributed by the bacteria or bacterial metabolites, as shown in murine model studies, is not fully investigated. Further investigations in human NKT cell at early development, along with the profile of gut metabolome is warranted.

## Acknowledgements

We thank J-I. Seo for technical assistance. This work was supported by the National Institutes of Health (K01-DK102771, R01-AT010268 and R01-AI165987: S.F.O.) and the National Research Foundation of Korea (2014R1A3A2030423 and 2012M3A9C4048780: S.B.P.; and Postdoctoral Fellowship Program: J-S. Y. and D-J.J.)

## Author contributions

S.F.O., D.L.K., S.B.P. and J-S.Y. conceived the idea and designed the outline of the research.

W.Z., N.G-Z., C.C.L., J-S.Y., M.W. and S.F.O. carried out LoF screening.

K.H. and B.G. carried out GoF screening.

J-S.Y., K.H. and B.G. carried out sphingolipid profiling of gut symbionts.

D-J.J., J-S.Y. and S.F.O. carried out *in vivo* NKT analysis.

J-S.Y. carried out *in silico* structural analysis, bacterial genetic manipulation, *in vitro* NKT activation assays and metagenomic analysis.

J-S.Y. and S.F.O. wrote the manuscript, and all authors contributed to the relevant discussion.

## Methods

### Mice

All animal procedures were supervised by the Harvard Center for Comparative Medicine and Brigham Women’s Hospital Center for Comparative Medicine and maintained by the institutional Animal Care and Use Facilities. Experimental groups were age-matched with one another. All mice are housed under 12-hour light-dark cycle and controlled climate (temperature: 21 ℃, humidity: 50%).

Germ-free Swiss-Webster mice were bred and maintained in inflatable plastic isolators. *B. fragilis* monocolonized mice were prepared by gavage of breeding pairs with a single bacterial strain (*B. fragilis* NCTC9343 wild-type, BF9343-Δ3069, BF9343-ΔagcT) and were maintained in isolators to obtain offspring (F1 and later generations). *E. faecalis* monocolonized mice were prepared by gavage of a pregnant female and were maintained in iso-P cages to obtain offspring. Stool samples from GF and monocolonized mice in isolators were regularly streaked onto plates and grown in both aerobic and anaerobic conditions to confirm sterility and absence of contamination. Specific pathogen–free (SPF) Swiss-Webster mice were purchased from Taconic. For *ex vivo* coculture assays conventional C57BL/6 mice were obtained from Taconic.

### Bacterial culture

Individual bacterial cultures, maintained as frozen stock, were first streaked onto plates; a single colony was picked up and inoculated into ∼1 mL of deoxygenated rich medium (2% proteose peptone, 0.5% yeast extract, 0.5% NaCl) supplemented with 0.5% D-glucose, 0.5% K_2_HPO_4_, 0.05% L-cysteine, 5 mg/L hemin and 2.5 mg/L vitamin K_1_) in an anaerobic chamber. Samples were grown overnight, centrifuged, and kept at –80°C until extraction. For Lactobacillus culture, MRS medium was used.

For bacterial plating, brain heart Infusion agar plates (3.7% brain heart infusion powder and 1.5% agar) supplemented with 5 mg/L hemin and 2.5 mg/L vitamin K_1_ was used. For selection, liquid media and agar plates were supplemented with the following antibiotics: 100 µg/mL ampicillin (*E. coli*), 10 µg/mL erythromycin (*B. fragilis*), 200 µg/mL gentamycin (*B. fragilis*) and 50 ng/mL anhydrotetracycline (*B. fragilis, B. thetaiotaomicron, and P. vulgatus*).

### Chemical compounds

Perdeuterated (d35) β-galactosylceramide (Matreya #1914), Sphinganine-d7 (d18:0) (Cayman #27145), N-omega-CD_3_-octadecanoyl-phytosphingosine (Matreya #2210), 3-keto sphinganine (d18:0) (Cayman #24380), βGluDAG (Avanti #840529), BbGL-2 (1-oleoyl-2-palmitoyl-3-(α-D-galactosyl)-sn-glycerol, Avanti #840528), MGlc-DAG (Avanti #840522p), KRN7000 (Avanti #867000) were obtained from commercial vendors.

### Lipidomic analyses

Perdeuterated (d35) beta-galactosylceramide (Matreya LLC), N-omega-CD_3_-Octadecanoyl-phytosphingosine (Matreya LLC), and Sphinganine-d_7_ (d18:0) (Cayman Chemical) as internal standards.

#### UHPLC-MS/MS condition

An UHPLC-MS/MS system (Thermo Scientific Vanquish RP-UPLC connected to a Q Exactive Orbitrap) was used for bacterial lipid profiling. Positive and negative ion mode method was established with parameters of spray voltage, 3.80 kV (positive ion mode) or 3.25 kV (negative ion mode); sheath gas, 40 AU; auxiliary gas, 8 AU; capillary temperature, 350°C; Aux gas heater temperature, 350°C; S-lens RF level, 65.0 AU; mean collision energy, 22.5 AU. YMC-Triart C8 column (2.1 mm × 100 mm × 3 µm, 200 µl min^−1^) was used for the gradient liquid chromatography elution (50% 2-propanol/10% acetonitrile/0.05% formic acid (0-2 mins), linearly increased to 85% 2-propanol/10% acetonitrile/0.05% formic acid (2-5 mins) and hold (5-12 mins), then returned to the initial condition for 0.1 min, and hold for 7.9 min), at 40°C. *Targeted lipidomics*. A high-resolution (R=70,000 @ m/z 200) MS1 scan (250–1000Da followed by top 3 data-dependent acquisition (R=17,500 @ m/z 200, isolation window: 1.0 m/z) to acquire MS and MS/MS spectra by Xcalibur 4.0 (Thermo Fisher Scientific). MS/MS spectra of biogenic and synthetic molecules were acquired and directly compared. Relative quantitation of individual bacterial lipids was done by quantitation of area under MS1 peaks with normalization by internal standard recovery.

### Transposon screening and targeted high-throughput LC-MS/MS lipidomics

Transposon insertion library of *B. fragilis* 638R were generated by conjugation with *E. coli* S17-lamda-pir harboring pSAM-Bt (Addgene_112497, deposited by Goodman and Gordon (Goodman et al., 2009)). Single colony of transposon-inserted mutants was isolated on BHI plate containing erythromycin 10 mg/mL and inoculated to 96-well plate. Individual samples were centrifuged and the pellets were dissolved and extracted with isopropyl alcohol. Supernatant were transferred to V-bottom 96 well plate, and analyzed by LC-MS/MS. An optimized, high-throughput method using Kinetex C8 (30mmx2.1mmx2.6um, column temperature 40 C) with 80% acetonitrile / 0.1% formic acid (0-0.1min), ramped to 85% 2-propanol / 0.1% formic acid (0.1-0.5 min) and hold (0.5-2.5 mins), then returned to the initial condition for 1.5min to re-equilibrate) was applied.

### In silico analysis

#### Metagenomic analysis

The species-level taxonomy profiles of multiple human gut metagenome datasets were obtained from curatedMetagenomicData v3.6.2 [10.1038/nmeth.4468]. Reference genomes of gut bacterial species within the profiles were downloaded using NCBI Entrez Direct and datasets tools. NCBI CD-search was used for domain search and DIAMOND [10.1038/s41592-021-01101-x] was used for protein blast search. Species contributions were analyzed and visualized by R with local polynomial regression fitting with shaded areas show 95% confidence intervals for the fit.

#### Structure superimposition

AlphaFold structures of AgcT (AF-A0A380YRQ3-F1) and BgsB (AF-Q830A6-F1) were obtained from UniProt database, the superimposition of AlphaFold structures of AgcT and BgsB were performed by ChimeraX matchmaker.

### RNA sequencing

Total RNA of *B. fragilis* NCTC 9343 and ΔBF9343_3069 were extracted using Trizol (ThermoFisher Scientific #15596026) and genomic DNA were removed by TURBO DNase (AM2238) treatment. The absence of gDNA contamination was confirmed by PCR of 16S rRNA DNA region. mRNA was enriched using a rRNA Depletion Kit (Bacteria) (NEB E7850). The sequencing library was constructed with Ultra™ II Directional RNA Library Prep with Sample Purification Beads (NEB E7765) and sequenced on sequencing machine. Paired-end reads were trimmed with. Trimmed reads were aligned to reference genome with Bowtie2. Read count matrix was obtained with featureCounts (v2.0.0). Gene counts were processed with DESeq2 (v1.32.0) to identify differentially expressed genes.

### Bacterial genetic manipulation for mutagenesis and heterologous expression

#### Bacteroides mutant and heterologous expression

*B. fragilis* NCTC 9343 mutants were constructed with a counterselection vector pLGB13, a gift from Laurie Comstock (Addgene 126618) [García-Bayona). Heterologous expression of *agcT* in *P. vulgatus* and *B. thetaiotaomicron* was conducted with a chromosome-integrated and inducible vector pNBU2 erm-TetR-P1T_DP-GH023 a gift from Andrew Goodman (Addgene 90324) [Lim 2017). Recombinant vectors were generated using HiFi DNA Assembly Cloning Kit (NEB E5520). During cloning procedures, Bacteroides were grown in brain heart infusion broth (3.7% brain heart infusion powder supplemented with five mg/L hemin and 2.5 mg/L vitamin K1) or brain heart infusion agar plates. Deletion or integration of the targeted locus were confirmed by PCR (Fig. S).

#### Enterococcus mutagenesis

*Enterococcus bgsB* mutants were generated using a counterselection vector pLT06(ref). Recombinant vectors were constructed using HiFi DNA Assembly Cloning Kit (NEB E5520). electroporation (device), single crossover selection on THB chloramphenicol plate, double crossover counter selection on THB phenylalanine plate.

### In vitro, APC-free NKT activity assay

24.7 NKT hybridoma cells were cultured in RPMI 1640 containing 10% FBS, sodium pyruvate, β-mercaptoethanol, NEAA at 37°C 5% CO2 (Vβ BV6S1 Dβ 2 Jβ 2.6 Vα AV14S1 Jα TRAJ15 281) [ref Behar 1999). Biotinylated murine CD1d monomers (NIH tetramer facility) were mixed with synthetic ligands or bacterial lipids in 50 mM pH 6.2 citrate buffer 0.25% CHAPS. After overnight incubation, ligand-loaded CD1d (0.25 ug per well) was bound onto Streptavidin-coated plates (R&D Systems #CP003). Plates were washed with PBS three times and then added 24.7 NKT cells (4 x 104∼105 per well) After overnight incubation, IL-2 of supernatants were analyzed by ELISA (BioLegend #431004).

### Colonic NKT cell analysis

#### Colonic lamina propria lymphocyte isolation

Conventional (SPF), GF and monocolonized mice were euthanized. The large intestines were collected and fat tissue was removed. The intestine was opened longitudinally, and fecal content was removed, cut into 1-inch pieces, and shaken in HBSS containing 2 mM EDTA for 50 min at 37°C. After the removal of epithelial cells, the intestines were washed in HBSS and incubated with RPMI 1640 containing 10% FBS, collagenase type VIII (1 mg/mL), and DNase I (0.1 mg/mL) (Sigma-Aldrich) for 45 min at 37°C under constant shaking.

The digested tissues were mixed with FACS buffer (PBS with 2% FBS and 1 mM EDTA), filtered twice through strainers (mesh size, 70 and 40 µm), and used for flow cytometry.

#### FACS analysis

For staining with the indicated dilution, APC-labeled mouse CD1d tetramer— unloaded or loaded with PBS-57 (1:500; NIH Tetramer Core Facility)—as well as anti-mouse CD3–FITC (1:400), TCRβ–PE (1:400), CD45–PerCP–Cy5.5 (1:200; Biolegend) and cell viability dye (Fixable Viability Dye eFluor™ 780, 1:1000; ThermoFisher) were used. Individual samples were stained for 20 min at 4°C and washed with cold FACS buffer. FACS analysis was performed with a BD FACSCanto system (BD Biosciences), pre-gated with forward and side scatters, a singlet population, and viable cells. The frequencies of CD3+/CD1d tetramer–positive cells from the gated total CD45+ population was enumerated as the target population. Data were analyzed and quantified with FlowJo V10 software (BD Biosciences).

### Statistical analysis

All data are shown as a represent of two or more independent experiment sets with the same trend. To enumerate significance, Adjusted P Value One-way ANOVA Dunnett’s multiple comparisons test is presented. *: P<0.05, **: P<0.01, ***: P<0.001.

## References

An, D., Na, C., Bielawski, J., Hannun, Y. A., & Kasper, D. L. (2011). Membrane sphingolipids as essential molecular signals for Bacteroides survival in the intestine. Proceedings of the National Academy of Sciences of the United States of America, 108 Suppl(Suppl 1), 4666–4671. https://doi.org/10.1073/pnas.1001501107

An, D., Oh, S. F., Olszak, T., Neves, J. F., Avci, F. Y., Erturk-Hasdemir, D., Lu, X., Zeissig, S., Blumberg, R. S., & Kasper, D. L. (2014). Sphingolipids from a Symbiotic Microbe Regulate Homeostasis of Host Intestinal Natural Killer T Cells. Cell, 156(1–2), 123–133. https://doi.org/10.1016/j.cell.2013.11.042

Brennan, P. J., Tatituri, R. V. V., Brigl, M., Kim, E. Y., Tuli, A., Sanderson, J. P., Gadola, S. D., Hsu, F. F., Besra, G. S., & Brenner, M. B. (2011). Invariant natural killer T cells recognize lipid self antigen induced by microbial danger signals. Nature Immunology 2011 12:12, 12(12), 1202– 1211. https://doi.org/10.1038/ni.2143

Brown, E. M., Ke, X., Hitchcock, D., Jeanfavre, S., Avila-Pacheco, J., Nakata, T., Arthur, T. D., Fornelos, N., Heim, C., Franzosa, E. A., Watson, N., Huttenhower, C., Haiser, H. J., Dillow, G., Graham, D. B., Finlay, B. B., Kostic, A. D., Porter, J. A., Vlamakis, H., … Xavier, R. J. (2019). Bacteroides-Derived Sphingolipids Are Critical for Maintaining Intestinal Homeostasis and Symbiosis. Cell Host & Microbe, 25(5), 668–680.e7. https://doi.org/10.1016/J.CHOM.2019.04.002

Cheng, T. Y., Relloso, M., Van Rhijn, I., Young, D. C., Besra, G. S., Briken, V., Zajonc, D. M., Wilson, I. A., Porcelli, S., & Moody, D. B. (2006). Role of lipid trimming and CD1 groove size in cellular antigen presentation. The EMBO Journal, 25(13), 2989–2999. https://doi.org/10.1038/SJ.EMBOJ.7601185

Comstock, L. E., Coyne, M. J., Tzianabos, A. O., Pantosti, A., Onderdonk, A. B., & Kasper, D. L. (1999). Analysis of a capsular polysaccharide biosynthesis locus of Bacteroides fragilis. Infection and Immunity, 67(7), 3525–3532. http://www.ncbi.nlm.nih.gov/pubmed/10377135

Coyne, M. J., Chatzidaki-Livanis, M., Paoletti, L. C., & Comstock, L. E. (2008). Role of glycan synthesis in colonization of the mammalian gut by the bacterial symbiont Bacteroides fragilis. Proceedings of the National Academy of Sciences of the United States of America, 105(35), 13099– 13104. https://doi.org/10.1073/PNAS.0804220105

Goodman, A. L., McNulty, N. P., Zhao, Y., Leip, D., Mitra, R. D., Lozupone, C. A., Knight, R., & Gordon, J. I. (2009). Identifying genetic determinants needed to establish a human gut symbiont in its habitat. Cell Host & Microbe, 6(3), 279–289. https://doi.org/10.1016/j.chom.2009.08.003

Johnson, E. L., Heaver, S. L., Waters, J. L., Kim, B. I., Bretin, A., Goodman, A. L., Gewirtz, A. T., Worgall, T. S., & Ley, R. E. (2020). Sphingolipids produced by gut bacteria enter host metabolic pathways impacting ceramide levels. Nature Communications, 11(1). https://doi.org/10.1038/S41467-020-16274-W

Kinjo, Y., Tupin, E., Wu, D., Fujio, M., Garcia-Navarro, R., Benhnia, M. R. E. I., Zajonc, D. M., Ben-Menachem, G., Ainge, G. D., Painter, G. F., Khurana, A., Hoebe, K., Behar, S. M., Beutler, B., Wilson, I. A., Tsuji, M., Sellati, T. J., Wong, C. H., & Kronenberg, M. (2006). Natural killer T cells recognize diacylglycerol antigens from pathogenic bacteria. Nature Immunology, 7(9), 978–986. https://doi.org/10.1038/ni1380

Le, H. H., Lee, M. T., Besler, K. R., & Johnson, E. L. (2022). Host hepatic metabolism is modulated by gut microbiota-derived sphingolipids. Cell Host & Microbe, 30(6), 798–808.e7. https://doi.org/10.1016/J.CHOM.2022.05.002

Lee, M. T., Le, H. H., Besler, K. R., & Johnson, E. L. (2022). Identification and characterization of 3-ketosphinganine reductase activity encoded at the BT_0972 locus in Bacteroides thetaiotaomicron. Journal of Lipid Research, 63(7). https://doi.org/10.1016/J.JLR.2022.100236

Liu, Q., Chan, A. K. N., Chang, W. H., Yang, L., Pokharel, S. P., Miyashita, K., Mattson, N., Xu, X., Li, M., Lu, W., Lin, R. J., Wang, S. Y., & Chen, C. W. (2022). 3-Ketodihydrosphingosine reductase maintains ER homeostasis and unfolded protein response in leukemia. Leukemia, 36(1), 100–110. https://doi.org/10.1038/S41375-021-01378-Z

Mayassi, T., Barreiro, L. B., Rossjohn, J., & Jabri, B. (2021). A multilayered immune system through the lens of unconventional T cells. Nature, 595(7868), 501. https://doi.org/10.1038/S41586-021-03578-0

Moremen, K. W., & Haltiwanger, R. S. (2019). Emerging structural insights into glycosyltransferase-mediated synthesis of glycans. Nature Chemical Biology 2019 15:9, 15(9), 853–864. https://doi.org/10.1038/s41589-019-0350-2

Naidenko, O. V., Maher, J. K., Ernst, W. A., Sakai, T., Modlin, R. L., & Kronenberg, M. (1999). Binding and Antigen Presentation of Ceramide-Containing Glycolipids by Soluble Mouse and Human Cd1d Molecules. Journal of Experimental Medicine, 190(8), 1069–1080. https://doi.org/10.1084/JEM.190.8.1069

Oh, S. F., Jung, D.-J., & Choi, E. (2022). Gut Microbiota-Derived Unconventional T Cell Ligands: Contribution to Host Immune Modulation. ImmunoHorizons, 6(7), 476–487. https://doi.org/10.4049/IMMUNOHORIZONS.2200006

Oh, S. F., Praveena, T., Song, H., Yoo, J. S., Jung, D. J., Erturk-Hasdemir, D., Hwang, Y. S., Lee, C. W. C., Le Nours, J., Kim, H., Lee, J., Blumberg, R. S., Rossjohn, J., Park, S. B., & Kasper, D. L. (2021). Host immunomodulatory lipids created by symbionts from dietary amino acids. Nature 2021 600:7888, 600(7888), 302–307. https://doi.org/10.1038/s41586-021-04083-0

Okino, N., Li, M., Qu, Q., Nakagawa, T., Hayashi, Y., Matsumoto, M., Ishibashi, Y., & Ito, M. (2020). Two bacterial glycosphingolipid synthases responsible for the synthesis of glucuronosylceramide and α-galactosylceramide. Journal of Biological Chemistry, 295(31), 10709–10725. https://doi.org/10.1074/jbc.ra120.013796

Olszak, T., An, D., Zeissig, S., Vera, M. P., Richter, J., Franke, A., Glickman, J. N., Siebert, R., Baron, R. M., Kasper, D. L. & Blumberg, R.S. (2012). Microbial Exposure During Early Life Has Persistent Effects on Natural Killer T Cell Function. Science, 336(6080), 489–493. https://doi.org/10.1126/science.1219328

Pasolli, E., Schiffer, L., Manghi, P., Renson, A., Obenchain, V., Truong, D. T., Beghini, F., Malik, F., Ramos, M., Dowd, J. B., Huttenhower, C., Morgan, M., Segata, N., & Waldron, L. (2017). Accessible, curated metagenomic data through ExperimentHub. Nature Methods 2017 14:11, 14(11), 1023–1024. https://doi.org/10.1038/nmeth.4468

Skelly, A. N., Sato, Y., Kearney, S., & Honda, K. (2019). Mining the microbiota for microbial and metabolite-based immunotherapies. In Nature Reviews Immunology (Vol. 19, Issue 5, pp. 305– 323). Nature Publishing Group. https://doi.org/10.1038/s41577-019-0144-5

Spears, M. E., Lee, N., Hwang, S., Park, S. J., Carlisle, A. E., Li, R., Doshi, M. B., Armando, A. M., Gao, J., Simin, K., Zhu, L. J., Greer, P. L., Quehenberger, O., Torres, E. M., & Kim, D. (2022). De novo sphingolipid biosynthesis necessitates detoxification in cancer cells. Cell Reports, 40(13). https://doi.org/10.1016/j.celrep.2022.111415

Stankeviciute, G., Tang, P., Ashley, B., Chamberlain, J. D., Hansen, M. E. B., Coleman, A., D’Emilia, R., Fu, L., Mohan, E. C., Nguyen, H., Guan, Z., Campopiano, D. J., & Klein, E. A. (2021). Convergent evolution of bacterial ceramide synthesis. Nature Chemical Biology 2021 18:3, 18(3), 305–312. https://doi.org/10.1038/s41589-021-00948-7

Stewart, C. J., Ajami, N. J., O’Brien, J. L., Hutchinson, D. S., Smith, D. P., Wong, M. C., Ross, M. C., Lloyd, R. E., Doddapaneni, H. V., Metcalf, G. A., Muzny, D., Gibbs, R. A., Vatanen, T., Huttenhower, C., Xavier, R. J., Rewers, M., Hagopian, W., Toppari, J., Ziegler, A. G., … Petrosino, J. F. (2018). Temporal development of the gut microbiome in early childhood from the TEDDY study. In Nature (Vol. 562, Issue 7728, pp. 583–588). Nature Publishing Group. https://doi.org/10.1038/s41586-018-0617-x

Theilacker, C., Sanchez-Carballo, P., Toma, I., Fabretti, F., Sava, I., Kropec, A., Holst, O., & Huebner, J. (2009). Glycolipids are involved in biofilm accumulation and prolonged bacteraemia in Enterococcus faecalis. Molecular Microbiology, 71(4), 1055–1069. https://doi.org/10.1111/J.1365-2958.2008.06587.X

Theilacker, C., Sava, I., Sanchez-Carballo, P., Bao, Y., Kropec, A., Grohmann, E., Holst, O., & Huebner, J. (2011). Deletion of the glycosyltransferase bgsB of Enterococcus faecalis leads to a complete loss of glycolipids from the cell membrane and to impaired biofilm formation. BMC Microbiology, 11(1), 1–11. https://doi.org/10.1186/1471-2180-11-67/TABLES/2

Zeissig, S., Olszak, T., Melum, E., & Blumberg, R. S. (2013). Analyzing antigen recognition by natural killer T cells. Methods in Molecular Biology, 960, 557–572. https://doi.org/10.1007/978-1-62703-218-6_41/COVER

