## Supplementary figures and images for "Functional and metagenomic level diversities of human gut symbiont-derived glycolipids"

### supplementary figures 1-6

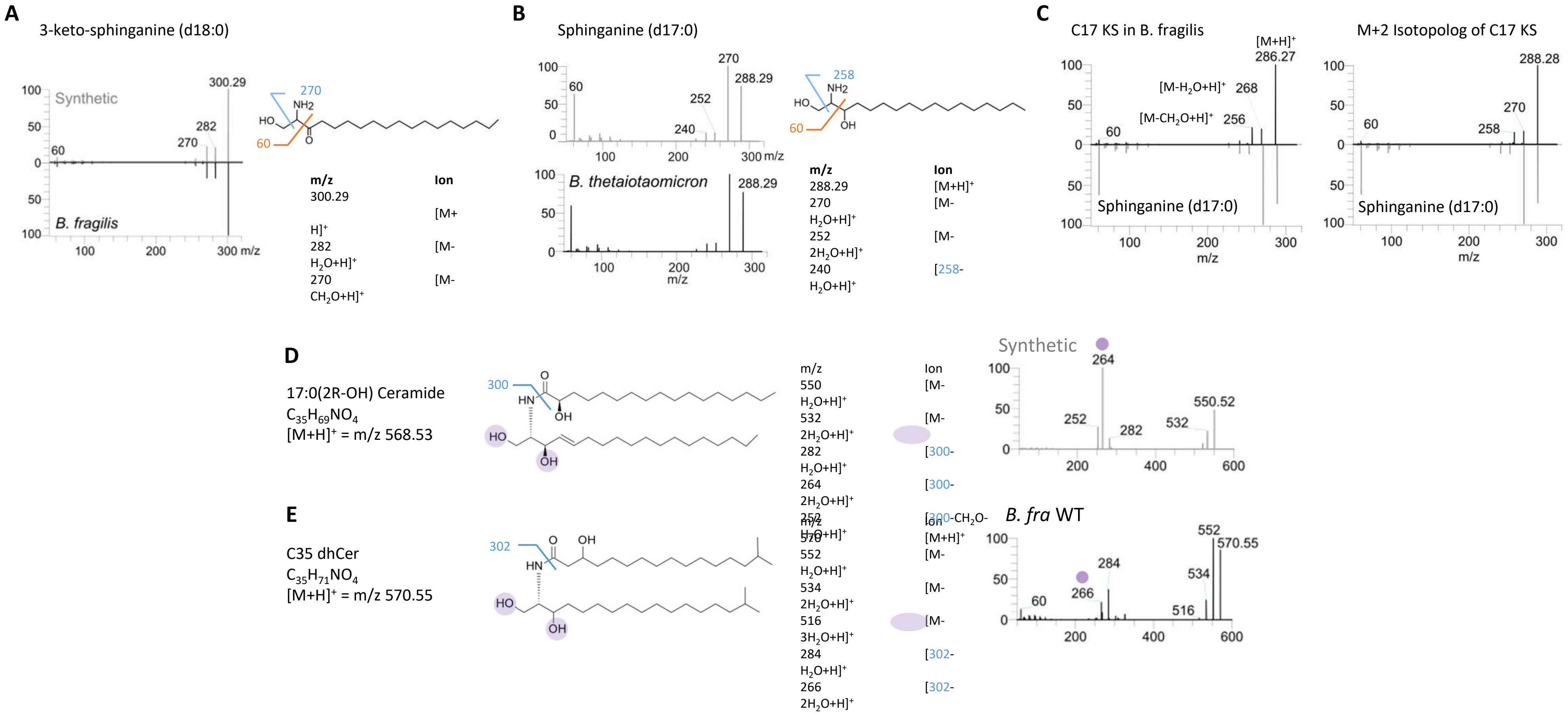

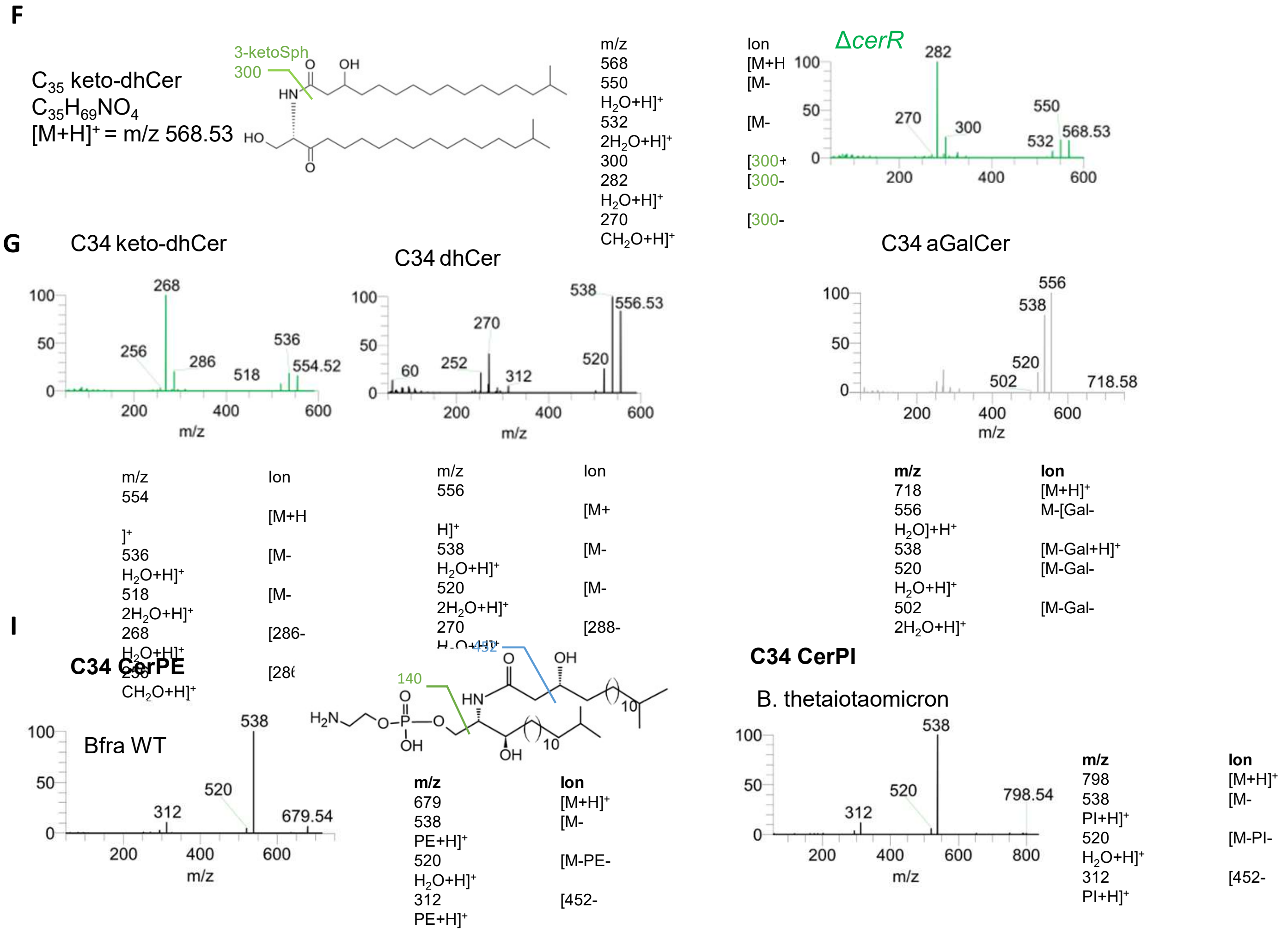


Figure S1 (cont'd)

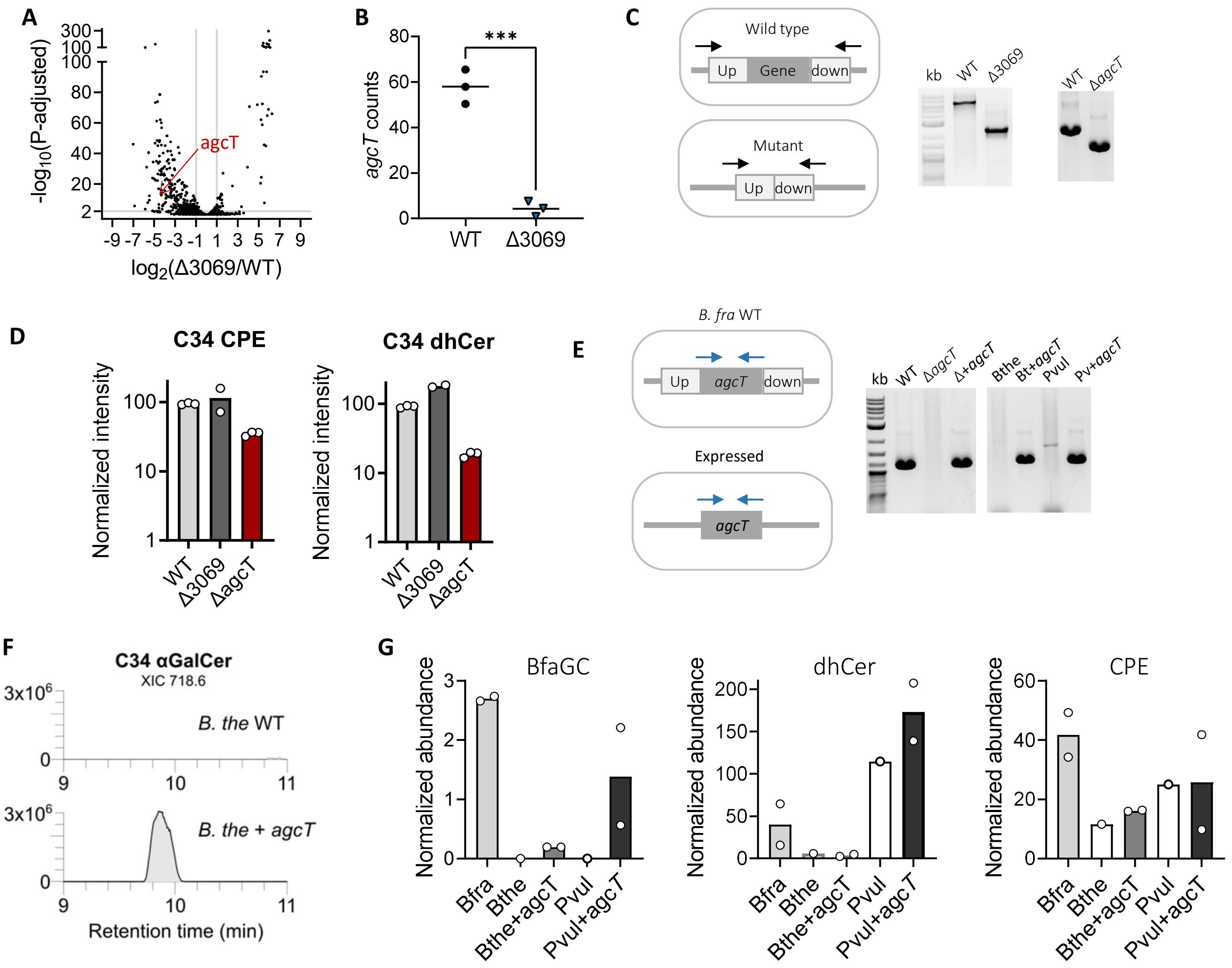

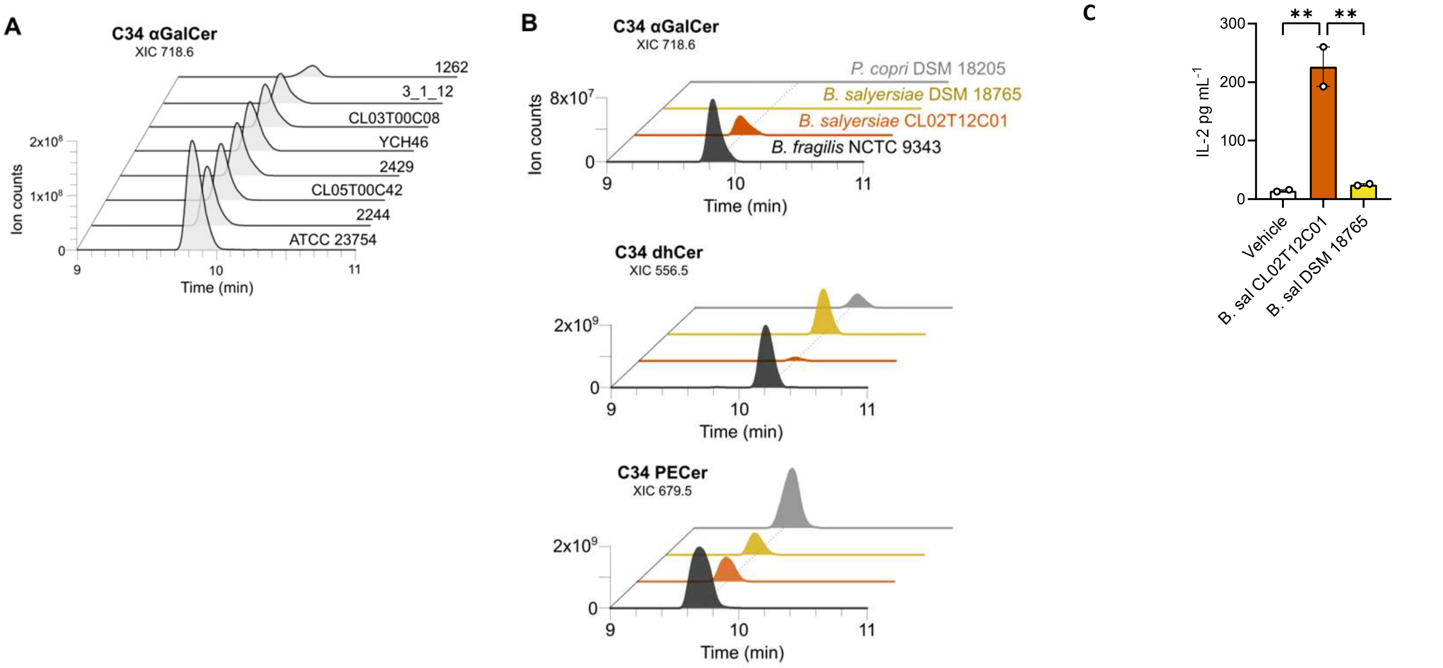

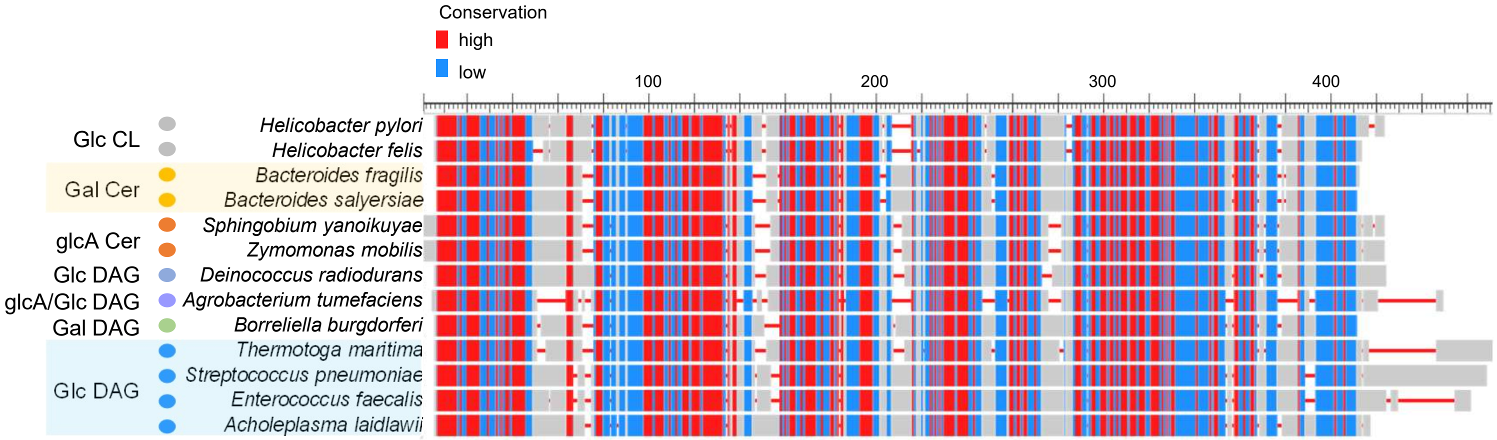

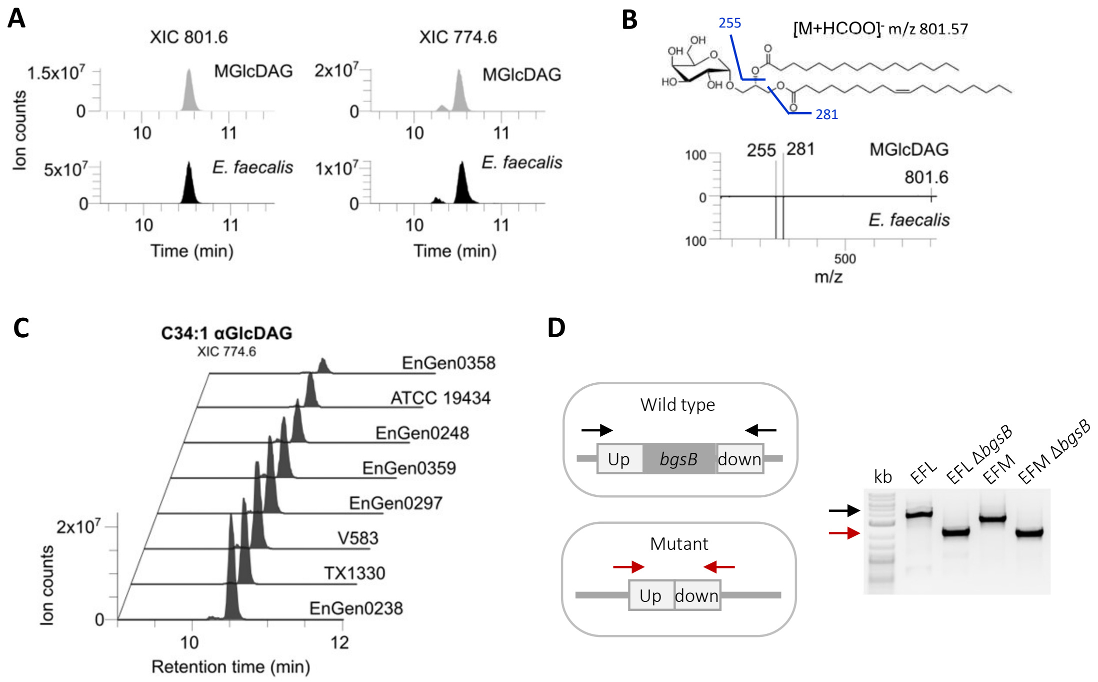

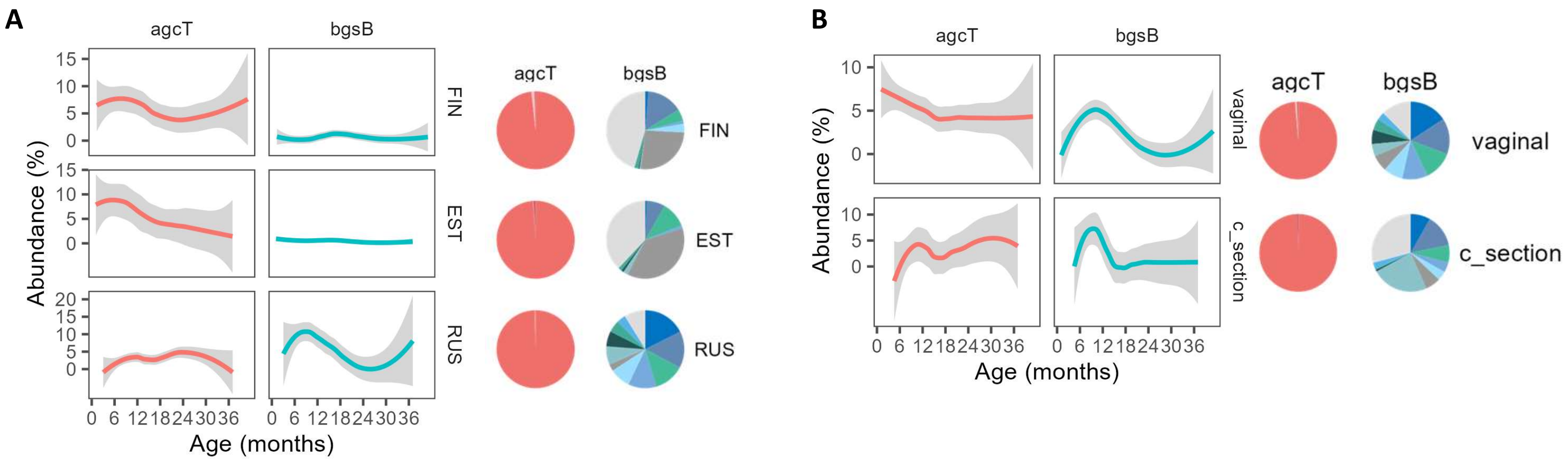
