## supplementary figure legends 1-6 for "Functional and metagenomic level diversities of human gut symbiont-derived glycolipids"

**Supplemental information**

**Supplementary figures 1-6**

**Supplementary figure legends**

**Figure S1. Tandem mass spectra analysis determined sphingolipid intermediate structures synthesized by Bacteroides.**

A) MS/MS spectra of 3-keto-sphinganines 3-KDS in B. fragilis. MS/MS spectral mirror plot of B. fragilis C18 3-KDS is identical to that of synthetic C18 3-KDS.

B) MS/MS spectra of Sphinganines in B. thetaiotaomicron. MS/MS spectra pattern of B. thetaiotaomicron C17 Sph matched that of synthetic C17 Sph. 240 m/z fragment ion represents the mass reduction of 18 Da, corresponding to the loss of a water molecule, from 258 m/z ion colored sky-blue.

C) MS/MS spectra of C17 KS in B. fragilis. KS is the major type of C17 sphingolipid molecule in B. fragilis. B. fragilis 288.28 m/z ion is isobaric to C17 Sph (B), but the MS/MS spectra pattern is different from that of C17 Sph. The ion fragmentation pattern of B. fragilis 288.28 m/z ion corresponds to the fragmentation pattern of an isotope of 286.27 m/z 3-KDS ion.

D) MS/MS spectra of C35 Cer synthetic compounds. The molecular ion at m/z=264, highlighted in a violet circle, can be generated by losing two water molecules from m/z 300 ion colored sky-blue.

E) MS/MS spectra of B. fragilis C35 dhCer. The molecular ion at m/z=266, highlighted in a violet circle, can be generated by the loss of two water molecules from 18:0 Sph at m/z=302, colored sky-blue.

F) MS/MS spectra of oxidized ceramides in ΔcerR. The molecular ion at m/z=568.53, isobaric with the synthetic C35 Cer ion in D, exhibits a distinct fragmentation pattern. The difference between a double-bond and hydroxyl residues can generate different fragmentation patterns. The fragment ion at m/z=300, colored light green, which does not have two hydroxyl groups, cannot generate the molecular ion at m/z=264, corresponding to the loss of two water molecules.

G) MS/MS spectra of C34 dhCer in B. fragilis WT and C34 oxCer in ΔcerR.

H) MS/MS spectral mirror plots of BfaGC exhibit exact matches to that of synthetic SB2217 compounds.

I) MS/MS spectra of B. fragilis C34 PE-Cer and B. thetaiotomicron C34 PI-Cer.

**Figure S2. Characterization of screened gene targets and generation of isogenic knockout and transformant strains.**

A) RNA-seq volcano plot for genes differentially expressed in Δ3069 compared to wild type.

B) The reduction of agcT expression in Δ3069 from the transcriptomic analysis. Unpaired two-tailed t-test.

C) Confirmation of isogenic KO strains of B. fragilis.

D) Relative abundance of C34 PE-Cer and BfaGC in wild type, Δ3069, and ΔagcT (ΔBF9343_3149).

E) Confirmation of agcT transformant strain in P. vugatus and B. thetaiotaimicron.

F) XIC of C34 BfaGC in B. thetaiotaomicron and AgcT-expressing B. thetaiotaomicron.

G) Relative abundance of C34 BfaGC, dhCer, and PE-Cer in two heterologously expressing strains.

**Figure S3. aGC production is limited to only a few gut symbionts.**

A) All *B. fragilis* strains, including type strains and clinical isolates, synthesize aGC.

B) *B. salyersiae* synthesizes small amount of aGC, with strain variability

C) *B. salyersiae* aGC can induce IL-2 from mouse NKT cells.

**Figure S4. Multiple alignments of cd03817 family proteins along with glycolipid products.**

**Figure S5. bgsB is responsible for Enterococcus aGlcDAG biosynthesis.**

A) Retention time of aGlcDAG of *E. faecalis* matches with commercially obtained monoglucosyl-DAG (MGlcDAG).

B) Mirror plot of MS/MS spectra of E. faecalis aGlcDAG vs MGlcDAG.

C) XIC of C34:1 aGlcDAG in various Enterococcus strains.

D) Confirmation of the generation of bgsB mutants in E. faecalis V583 and E. faecium ATCC 19434.

**Figure S6. Gut metagenomic landscapes of agcT and bgsB signature-bearing species during early age.**

A-B) agcT+ profile and bgsB+ profile during early age subset by A) country and B) delivery mode.
